# CA3 place cells that represent a novel waking experience are preferentially reactivated during sharp wave-ripples in subsequent sleep

**DOI:** 10.1101/398560

**Authors:** Ernie Hwaun, Laura Lee Colgin

## Abstract

A popular model of memory consolidation posits that recent memories stored in the hippocampus are reactivated during sleep and thereby transferred to neocortex for long-term storage. This process is thought to occur during sharp wave-ripples (SWRs) in non-rapid eye movement (NREM) sleep. But whether the hippocampus consolidates all recent memories in the same manner remains unclear. An efficient memory system may extract novel information from recent experiences for preferential consolidation. In the hippocampus, memories are thought to be stored initially in CA3. Therefore, CA3 place cells that encode novel experiences may be preferentially reactivated during SWRs in subsequent sleep. To test this hypothesis, we recorded CA3 place cells in rats during exposure to a familiar and a novel environment and during subsequent overnight sleep. We found that cells that preferentially coded a novel environment showed larger firing rate increases during SWRs in NREM sleep than cells that preferentially coded a familiar environment. Moreover, CA3 place cell ensembles replayed trajectories from a novel environment during NREM sleep with higher fidelity than trajectories from a familiar environment. Together, these results suggest that CA3 representations of novel experiences are preferentially processed during subsequent sleep.

## Introduction

Damage to the hippocampus impairs the ability to form new episodic memories while sparing the ability to recall remote memories (Scoville and Milner, 1957; Squire, 1992; Squire et al., 2004). This selective memory impairment inspired the proposal that recent memories are initially stored in the hippocampus and then later consolidated in neocortex for long-term storage (McClelland et al., 1995). This memory consolidation process is thought to occur during sleep (Diekelmann and Born, 2010). Recordings of hippocampal “place cells”, neurons that exhibit spatially selective firing (O’Keefe and Dostrovsky, 1971; O’Keefe, 1976), have provided support for the idea that hippocampal memories are consolidated during sleep. Place cell ensemble activity during non-rapid eye movement (NREM) sleep has been shown to reflect activity from previous awake experiences (Wilson and McNaughton, 1994; Skaggs et al., 1996; Kudrimoti et al., 1999; Nádasdy et al., 1999; Lee and Wilson, 2002), and such recurring ensemble activity has thus been termed “replay”. Replay of hippocampal firing patterns during NREM sleep tends to occur during sharp wave-ripples (SWRs) (Nádasdy et al., 1999; Lee and Wilson, 2002), the dominant hippocampal network pattern during NREM sleep (Buzsáki, 1986). Brief disruption of hippocampal activity after the onset of SWRs during post-learning sleep and rest has been reported to impair spatial memory retrieval (Girardeau et al., 2009; Ego-Stengel and Wilson, 2010), providing causal evidence that SWR-related replay supports memory consolidation.

However, the hippocampus may not consolidate all recent memories equally. Memories of novel experiences may be preferentially reactivated and consolidated, perhaps to reduce redundancy in memory storage. Indeed, CA1 place cells that represent novel experiences have been shown to be preferentially reactivated during ripples and high frequency ripple-like events (Cheng and Frank, 2008; O’Neill et al., 2008). However, memory consolidation requires output from hippocampal subregion CA3 (Nakashiba et al., 2009), and most studies have only investigated replay in CA1 place cells (Lee and Wilson, 2002; Foster and Wilson, 2006; Csicsvari et al., 2007; Diba and Buzsáki, 2007), have grouped CA1 and CA3 place cells together (Karlsson and Frank, 2009), or have recorded too few CA3 cells to compare novel and familiar conditions (O’Neill et al., 2008). Place cell ensemble representations of novel environments develop more slowly in CA3 than in CA1 (Leutgeb et al., 2004), and thus it is unclear whether place cell representations of novel environments are processed in CA3 in the same manner as in CA1. Thus, it remains unclear whether CA3 place cells representing novel experiences are selectively reactivated during SWRs.

Moreover, decreases in firing rates during REM compared to NREM have been reported previously for CA1 and CA3 (Grosmark et al., 2012; Mizuseki et al., 2012). These firing rate decreases during REM may reflect homeostatic processes that reset hippocampal activity levels after memories have been consolidated. However, it remains unclear whether such firing rate decreases during REM occur preferentially for cells encoding novel experiences.

To address the above questions, we recorded place cells in area CA3 of rats navigating both familiar and novel environments and during subsequent overnight sleep. We compared firing patterns of CA3 place cells that coded familiar and novel locations to assess whether novelty-dependent activity changes were observed during NREM and REM sleep. Moreover, we investigated whether replay events from CA3 place cell ensembles during NREM sleep preferentially reflected novel trajectories, as expected if replay in CA3 plays a role in consolidation of newly formed memories. The results suggest that CA3 place cells that code novel experiences are preferentially recruited during SWRs and display higher fidelity replay than CA3 place cells that code familiar experiences.

## Materials & Methods

### Subjects

Five male Long-Evans rats weighing from ∼400 to 600 g were used in this study. The age of the rats ranged from 20 weeks to 60 weeks old at the time of behavioral testing. Data from four of these rats were included in a previous study (Trettel et al., 2017). Rats were housed in custom-built acrylic cages (40cm x 40cm x 40cm) on a reverse light cycle (Light: 8pm to 8am). The cages contained enrichment materials (e.g., plastic balls, cardboard tubes, and wooden blocks). Active waking behavior recordings were performed during the dark phase of the cycle, and overnight sleep recordings were performed during the light phase of the cycle (i.e., from 8 pm to 8 am). Rats recovered from surgery for at least one week prior to the start of behavioral testing. During the data collection period, rats were placed on a food-deprivation regimen that maintained them at no less than ∼90% of their free-feeding body weight. All experiments were conducted according to the guidelines of the United States National Institutes of Health Guide for the Care and Use of Laboratory Animals under a protocol approved by the University of Texas at Austin Institutional Animal Care and Use Committee.

### Surgery and tetrode positioning

Recording drives with 14 independently movable tetrodes were surgically implanted above the right hippocampus (anterior-posterior [AP] 3.8 mm, medial-lateral [ML] 3.0 mm, dorsal-ventral [DV] 1 mm). Bone screws were placed in the skull, and the screws and the base of the drive were covered with dental cement to affix the drive to the skull. Two screws in the skull were connected to the recording drive ground. Before surgery, tetrodes were built from 17 μm polyimide-coated platinum-iridium (90/10%) wire (California Fine Wire, Grover Beach, California). The tips of tetrodes designated for single unit recording were plated with platinum to reduce single channel impedances to ∼150 to 300 kOhms. Over the course of ∼1 month following surgery, tetrodes were gradually lowered into the hippocampus (1-4 tetrodes targeting CA1 with final DV depth of ∼2 mm, and 8-11 tetrodes targeting CA3 with final DV depth of ∼3 mm for each rat). One tetrode was designated as the reference for differential recording and remained in a quiet area of the cortex above the hippocampus throughout the experiments. This tetrode was moved up and down until a quiet location was found and was continuously recorded against ground to ensure that it remained quiet throughout data collection. Another tetrode was placed in the CA1 apical dendritic layers to monitor local field potentials (LFPs) in the hippocampus during placement of the other tetrodes.

### Data acquisition

Data were acquired using a Digital Lynx system and Cheetah recording software (Neuralynx, Bozeman, Montana). The recording setup has been described in detail previously (Hsiao et al., 2016; Zheng et al., 2016). Briefly, LFPs from one randomly chosen channel within each tetrode were continuously recorded in the 0.1-500 Hz band at a 2000 Hz sampling rate. Input amplitude ranges were adjusted before each recording session to maximize resolution without signal saturation. Input ranges for LFPs generally fell within +/-2000 to +/-3000 μV.

To detect unit activity, signals from each channel in a tetrode were bandpass filtered from 600 to 6000 Hz. Spikes were detected when the signal on one or more of the channels exceeded a threshold set daily by the experimenter, which ranged from 55-65 μV. Detected events were acquired with a 32000 Hz sampling rate for 1 ms.

Signals were recorded differentially against a dedicated reference channel (see “Surgery and tetrode positioning” section above). Video was recorded through the Neuralynx system with a resolution of 720 x 480 pixels and a frame rate of 29.97 frames per second. Animal position and head direction were tracked via an array of red and green light-emitting diodes (LEDs) on an HS-54 headstage (Neuralynx, Bozeman, Montana) attached to a recording drive.

### Behaviors and overnight sleep

Rats were trained to run unidirectionally on a circular track (diameter of 100 cm, height of 50 cm, and width of 11 cm) to retrieve food rewards (i.e., pieces of Froot Loops) at opposite ends of the track for at least three days in a familiar room. Four ten-minute sessions were conducted daily, separated by ten-minute rest sessions. Once tetrodes reached their target recording locations (i.e., CA3 and CA1), rats performed the circular track task in the afternoon (from 6 pm to 8 pm), and then were placed in a square box (60 cm x 60 cm with 50 cm high walls) in the familiar room for overnight sleep recording (from 8 pm to 8 am). On the next day, rats ran the same circular track task in the familiar room in the first and fourth sessions. However, on this day, the rats ran on a different circular track in a novel room (i.e., a room where the rats had not been previously) during the second and third sessions (Figure 1A). Small raised circles of various colors were adhered to the novel circular track to make its texture and appearance distinct from that of the familiar circular track in the familiar room. However, the dimensions of both novel and familiar circular tracks were the same. Upon completion of the circular track task, the rats again stayed in the square box in the familiar room for overnight recording (again from 8 pm to 8 am). Putative sleep episodes were identified by finding extended periods of immobility (>=60 seconds) in the box using custom code written in Matlab (MathWorks). Only sleep episodes that were also visually confirmed by two independent researchers were included for further analysis. The same behavioral protocol including recordings in the familiar and novel rooms was repeated the next morning (from 9 am to 11 am) to check whether cell ensemble recordings remained stable overnight.

**Figure 1.**
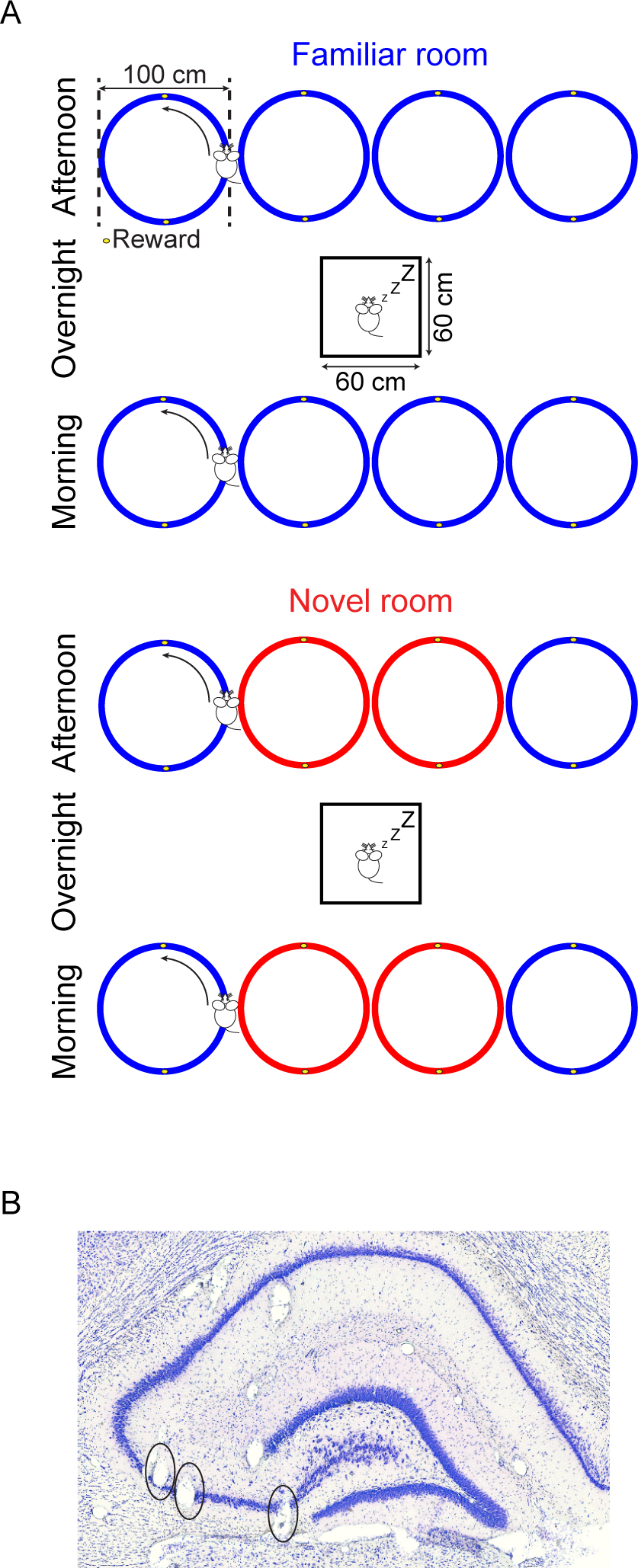
Behavioral protocol and verification of recording locations. ***A***, A schematic of the experimental setup shows the sequence of behavior recordings. Rats were trained to run unidirectionally on a circular track to retrieve food rewards in a familiar room (in blue) for four sessions. After completing the circular track task, rats were placed in a square box for recordings of overnight sleep. On the next morning, the same behavioral protocol was repeated. In the afternoon on the same day, the same behavioral training protocol was repeated except the middle two sessions took place on a novel track in a novel room (in red; this color scheme is used consistently throughout the paper). These behavioral sessions were again followed by overnight sleep recording. The same behavioral training sequence was then repeated the next morning to check whether place cells remained stable across overnight recordings. Note the novel room in the morning was the same novel room as the previous afternoon. ***B***, Example histologically verified CA3 recording sites are shown (indicated by black ovals).

### Identifying periods of REM and NREM sleep

Periods of REM sleep were detected offline using the ratio of the theta band (6-10 Hz) to delta band (2-5 Hz) wave amplitude (Csicsvari et al., 1999) using LFPs recorded from a tetrode placed in the CA1 apical dendritic layers. The ratio was smoothed by a moving average time window of 5 seconds. Sleep periods with a ratio greater than 2 were categorized as REM. Any successively detected periods with an intervening time interval less than 3 seconds were combined. Detected REM periods with a duration of less than 10 seconds were excluded. Detected REM sleep was verified by visually determining that there was a sustained increase in theta band amplitude in time-resolved amplitude spectrograms, and boundaries were adjusted manually if necessary. The spectrogram was estimated using a complex Morlet wavelet transform with a width parameter of 6 periods. The remainder of identified sleep periods that were not classified as REM were categorized as NREM. NREM periods with a duration of less than 5 seconds were excluded.

### Identifying SWRs in NREM sleep

LFPs recorded from tetrodes with CA1 units were used to detect SWRs in NREM sleep. The detection method used in this study has been described in detail previously (Cheng and Frank, 2008). Briefly, LFPs were first bandpass filtered between 150 and 250 Hz. A Hilbert transform was implemented to estimate the instantaneous wave amplitude of the filtered LFPs by taking the absolute value of the complex signal. The amplitude was then smoothed with a Gaussian kernel with a standard deviation of 25 ms. A SWR event was detected if the amplitude exceeded 3 standard deviations above the mean for at least 15 ms. Detected SWR events were bounded by first crossings of the mean amplitude. Overlapping SWR events were combined across CA1 tetrodes, so events could extend beyond a SWR detected on a single tetrode (Karlsson and Frank, 2009). Any SWR event detected during identified REM periods was excluded from further analysis (2169 out of 53919, or ∼4%, detected events were excluded).

### Spike sorting and unit classification

We performed spike sorting offline using graphical cluster-cutting software (MClust; A.D. Redish, University of Minnesota, Minneapolis). Spikes were clustered manually using two-dimensional projections of three different features of spike waveforms (i.e., energies, peaks, and peak-to-valley differences) from four channels. We excluded putative fast-spiking interneurons (units with mean firing rates > 5 Hz). We measured the quality of clusters using L ratio (0.294 ± 0.0271) and isolation distance (38.2 ± 3.67) measures (Schmitzer-Torbert et al., 2005). The two measures were calculated using energies, peaks, and peak-to-valley differences on each channel. We categorized single units into four different groups based on whether they were active only in the novel room (Novel), only in the familiar room (Familiar), in both rooms (Both), or in neither room but active during sleep (Neither). No differences in measures of cluster quality were observed across the different groups (Supplementary Figure 1). Because we were unable to determine whether units in the “Neither” category were place cells with place fields in another environment (e.g., an area of their home cage) or a different type of cell, units in the Neither category were not analyzed further.

A unit was considered to be active during waking behaviors if it exhibited a peak firing rate of at least 1 Hz (see “Unit firing analysis*”* section below). Units that were not active during waking behaviors, but had valid clusters from sorting of spikes during sleep, were considered active during sleep. We also quantified how selective a unit was for the novel versus the familiar room using a selectivity index. The selectivity index was computed using the following formula: (μ_novel_ – μ_familiar_) / (μ_novel_ + μ_familiar_), where μ was the mean firing rate in either the familiar or novel room. The selectivity index ranged from −1 to 1, with −1 indicating that a unit was exclusively active in the familiar room and 1 indicating that a unit was exclusively active in the novel room.

### Unit firing analysis

We calculated one-dimensional position tuning for each unit by binning the spikes into 5 degree bins and dividing the number of spikes in each bin by the amount of time spent in each bin. We estimated the position tuning for each lap on the circular track. To exclude out-of-field spikes during the immobile state, we only included times when rats were moving at least 5 cm/s (or equivalently 5.7 degrees/s) to estimate position tuning. We then smoothed the position tuning curves through convolution with a Gaussian kernel (SD = 5 degrees). Units with peak firing rates of at least 1 Hz on the circular track in either room were considered active during waking behavior (as described above).

We constructed a population vector containing firing rates of all units for each position bin in each lap from the normalized position tunings of all simultaneously recorded active units (Leutgeb et al., 2005). The position tuning for each unit across laps was normalized by dividing each unit’s maximal firing rate across laps. The population vectors for each position bin within each lap were further concatenated into a lap vector. We computed the Pearson’s correlation coefficient between pairs of lap vectors to assess how similar population activities from a pair of laps were as a function of lap number difference. A larger lap number difference indicated that a pair of laps were further apart in time. We computed population vector correlations separately for pairs of laps in novel and familiar environments (Figure 2B).

**Figure 2.**
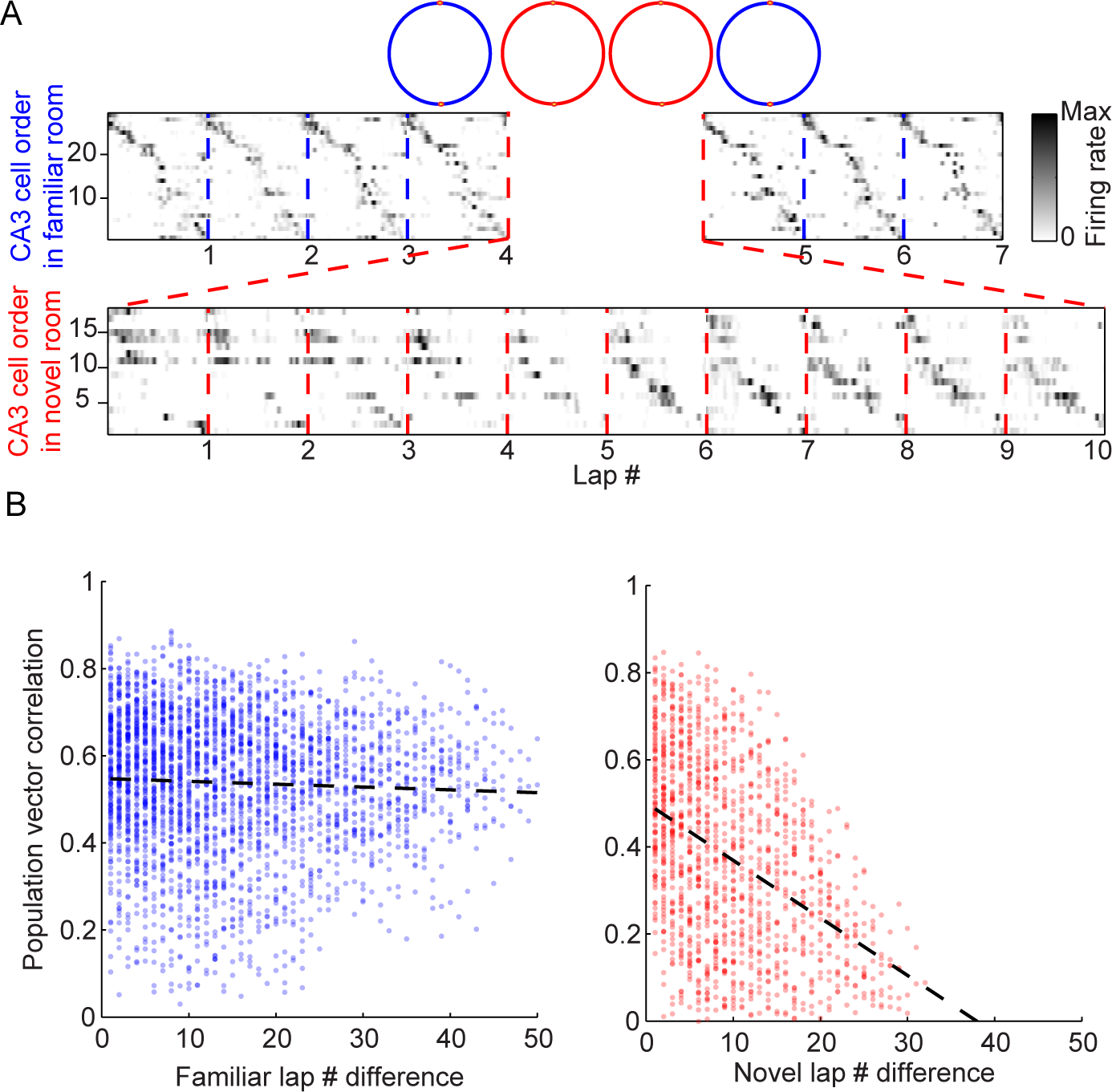
CA3 representations of a novel track develop across laps. ***A***, Examples of simultaneously recorded CA3 place cells on the circular track in the familiar room (blue) and in the novel room (red) are shown. Vertical dashed lines separate each lap on the track. Angular positions for each lap on the circular track were unwrapped on the x-axis (shown for each lap in between lap numbers). CA3 place cells active in either the familiar or the novel room were sorted on the y-axis according to their peak firing positions. Note that ensembles of CA3 place cells active in the familiar and novel rooms were different. Grayscale indicates normalized firing rate, with the darkest color representing maximal firing rate. In this example, most of the CA3 place cells that were active on the familiar track fired consistently at a specific position along the track, as shown by a roughly diagonal pattern of place cell ensemble activity that appears stable across laps. In contrast, most of the CA3 place cells that were active on the novel track changed their firing locations and/or firing rates across laps. ***B***, CA3 ensemble activity across laps was more similar in the familiar room than in the novel room. Population vector correlations were employed to quantify similarity between place cell ensemble representations for pairs of laps (see Materials and Methods). Lap number differences reflected how far apart pairs of laps were in time (e.g., laps 1 and 10 would have a lap number difference of 9), and a pair of laps were further apart in time if their lap number difference was higher. Each dot represents a correlation value for a pair of laps in the same room. Black dashed lines show the linear fits to the data.

To examine unit firing during REM sleep, we normalized the duration of each sleep episode (i.e. 100% corresponds to entire duration of each episode) that contained REM periods to obtain a mean firing rate across time for each unit (Grosmark et al., 2012). Each normalized REM and NREM period was divided into equal thirds. Since individual units had different baseline firing rates, we additionally normalized the firing rates by dividing by the mean firing rate during the NREM periods that preceded the REM periods to facilitate comparisons between different groups of units (see “Spike sorting and unit classification*”* section above).

To investigate unit firing on a finer time scale during NREM sleep, we aligned the onset of all detected SWRs and averaged firing rates across all SWRs for each unit. We obtained the firing rate around SWRs by binning the spikes into 1 ms time bins. We smoothed the firing rate with a Gaussian kernel (SD = 5 ms). To account for baseline firing rate differences, we normalized the firing rates in two different ways. The first way (Figure 3) was to divide the firing rate of each CA3 place cell by its associated mean firing rate during the period before ripple onset (i.e., from 0.4 to 0.1 seconds prior to ripple onset). The second way (Supplementary Figure 2) was to divide the firing rate of each CA3 place cell by its mean firing rate during SWRs in NREM sleep recorded the night before the novel experience.

**Figure 3.**
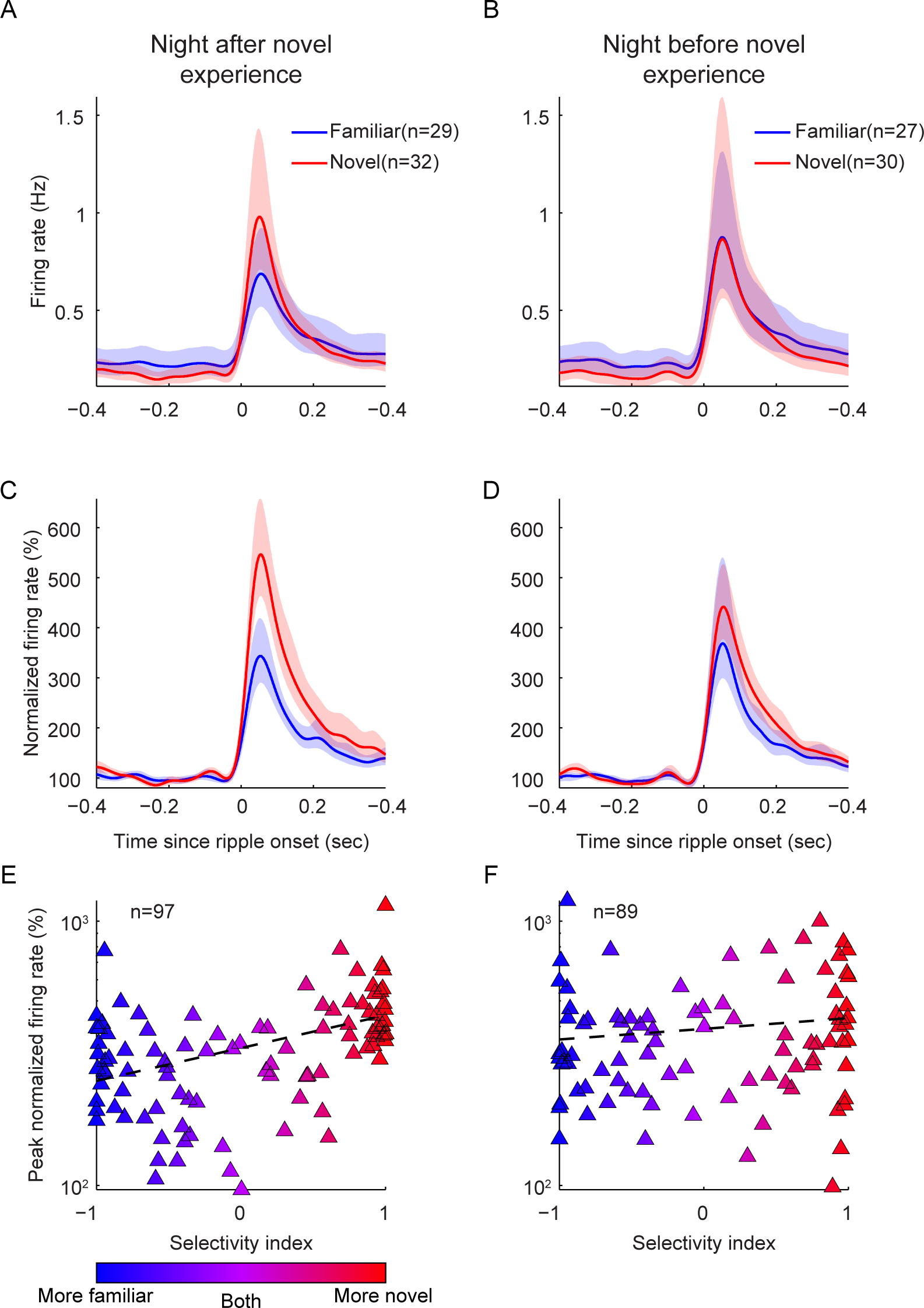
CA3 place cells that preferentially represented a novel environment showed stronger increases in firing rates during SWRs. ***A-B***, Mean firing rates in CA3 increased after the onset of SWRs during sleep following (A) and preceding (B) a novel experience. Firing rates for each CA3 place cell were averaged across detected SWRs. Shaded areas represent 95% confidence intervals. ***C-D***, Normalized firing rates of CA3 place cells in the Novel group increased more after the onset of SWRs than CA3 place cells in the Familiar group for sleep following (C) but not preceding (D) a novel experience. Mean firing rates near SWRs were normalized by the mean firing rates during the period before ripple onset (i.e., from 0.4 to 0.1 seconds prior to ripple onset). ***E-F***, Peak normalized firing rates during SWRs were positively correlated with a selectivity index during sleep following (E) but not preceding (F) a novel experience. The selectivity index quantified how selective activity of a CA3 place cell was for the familiar room (selectivity index = −1 for maximal selectivity for the familiar room) or the novel room (selectivity index = 1 for maximal selectivity for the novel room). Both color scale and horizontal position indicate the degree of selectivity. The black dashed line shows the linear fit to the data.

### Decoding accuracy analysis

We used a Bayesian decoding algorithm (Zhang et al., 1998) to compute the probability of a rat’s angular position given ensemble activity of recorded CA3 place cells. The position with the highest probability was taken as the decoded position. To ensure stable spatial representations across overnight sleep, we used the position tunings of each CA3 place cell during active behaviors on the circular track before sleep to decode rats’ positions during active behaviors on the circular track after sleep. Positions were decoded across 500 ms time windows using a time step of 100 ms. Only time windows when rats were running more than 5 cm/s were included.

We quantified decoding accuracy measures for each rat using two different measures that have been previously described (Davidson et al., 2009). First, decoded probability distributions were averaged across every pass through each position on the track (Figure 4B & 4C, top panels). For each rat, 60% of the total probability density was required to be less than 30 degrees away from the actual position to be included for further analyses (Rat 100’s familiar session and Rat 140’s familiar and novel sessions were excluded based on this criterion). The second measure took the differences between decoded positions and actual positions as errors, and cumulative distributions of errors were determined (Figure 4B & 4C, bottom panels). Only rats with an error distribution reaching 60% at error values less than 30 degrees were included for further analysis (Rat 140’s familiar and novel sessions did not meet this criterion but had already been excluded-see above).

**Figure 4.**
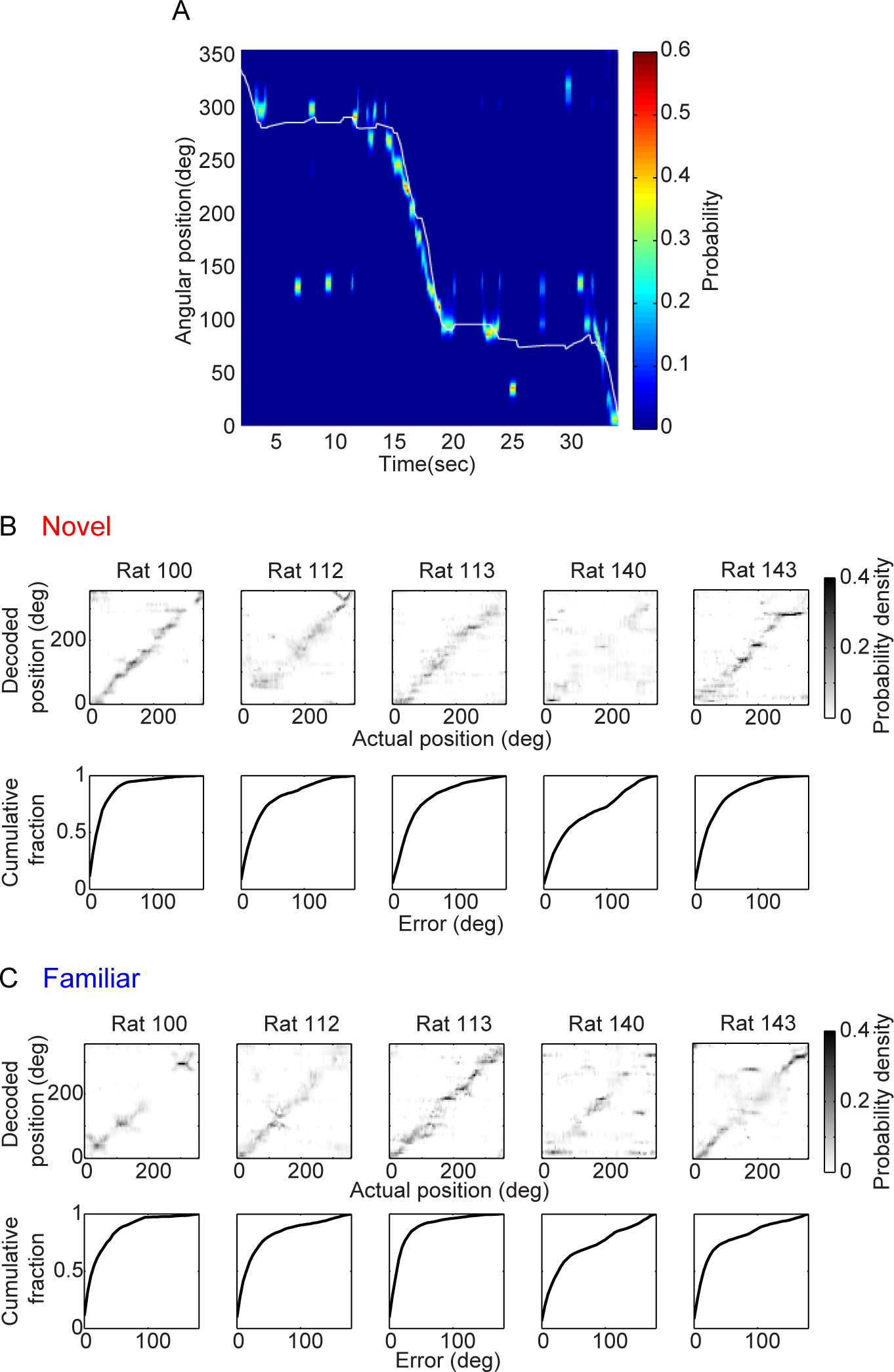
Decoding accuracy across active waking behaviors. ***A***, An example Bayesian decoded spatial probability distribution for a CA3 place cell ensemble recorded on the circular track shows that a rat’s actual position could be decoded accurately, especially during active movement. The color scale indicates the decoded probability distribution of angular position, with warmer colors representing higher probability. The white line shows the actual position of the rat. ***B-C***, (Top) Confusion matrices show mean decoded probability distributions across rats’ actual positions at times when movement speeds on the circular track > 5 cm/s. A continuous diagonal stripe in a confusion matrix corresponds to accurate decoding. (Bottom) Cumulative distributions of decoding errors are presented for each rat. Decoding errors were defined as the difference between a rat’s actual position at a given time and the position with the highest decoded probability at that time. Panel B shows data from the novel room, and panel C shows data from the familiar room.

### Sleep replay analysis

We defined candidate replay events during NREM sleep as the periods of time when the population firing rate of a subset of CA3 place cells exceeded 3 standard deviations above the mean, bounded by first crossings of the mean (Pfeiffer and Foster, 2013). Only candidate events with a minimum of 5 different active cells and a duration ranging from 50 to 500 ms were included for further analysis. According to whether a given subset of CA3 place cells was active in the novel or the familiar room (i.e. peak firing rate >= 1 Hz, see above), candidate events were categorized into Novel or Familiar groups, respectively. Since some CA3 cells were active in both rooms, some of the candidate events in Novel and Familiar groups overlapped with each other in time (53 out of 244 and 53 out of 120 events in Novel and Familiar groups, respectively). To avoid double counting, candidate events that overlapped in time were excluded from further analysis.

To evaluate whether a familiar or a novel experience was replayed during NREM sleep, we applied the Bayesian decoding method described above to successive 20 ms time windows within a candidate replay event using a time step of 10 ms. A standard method was used to assess the fidelity of replay, which involved fitting a line to decoded distributions across time and then using associated r^2^ values as a measure of replay fidelity (Davidson et al., 2009; Karlsson and Frank, 2009). r^2^ values range from zero to one, with values closer to one corresponding to higher fidelity replay. Since the behavioral apparatus was a circular track, we employed circular-linear regression to obtain r^2^ values (Kempter et al., 2012).

### Histology

At the end of the experiments, rats were sacrificed by injecting (i.p.) a lethal dose of pentobarbital. Rats then received intra-cardiac perfusion with phosphate-buffered saline followed by formalin. Brains were then sliced into 30 µm coronal sections and stained with cresyl-violet to confirm final tetrode positions in CA3 (Figure 1B).

### Experimental design and statistical analysis

Due to a limited number of recording rooms available at the time of data collection, we were only able to expose most of the rats to one novel room (except Rat 143 who was exposed to two different novel rooms). Rats were exposed to the novel room on the recording day when we estimated that our yield of simultaneously recorded CA3 place cells was maximal. The number of simultaneously recorded CA3 cells for each recording session is reported in Table 1. Note that the set of CA3 cells recorded in the second novel room for Rat 143 were different than the CA3 cells recorded in the first novel room. In Table 2, the number of CA3 cells is further divided into different cell groups for each overnight recording session following exposure to a novel room. In all five rats, multiple CA3 place cells were included in all four groups (i.e., Novel, Familiar, Neither, Both; Table 2). However, CA1 place cells were included in both Novel and Familiar groups in only two rats (Table 3). Thus, CA1 place cells were not included in this study.

**Table 1.**
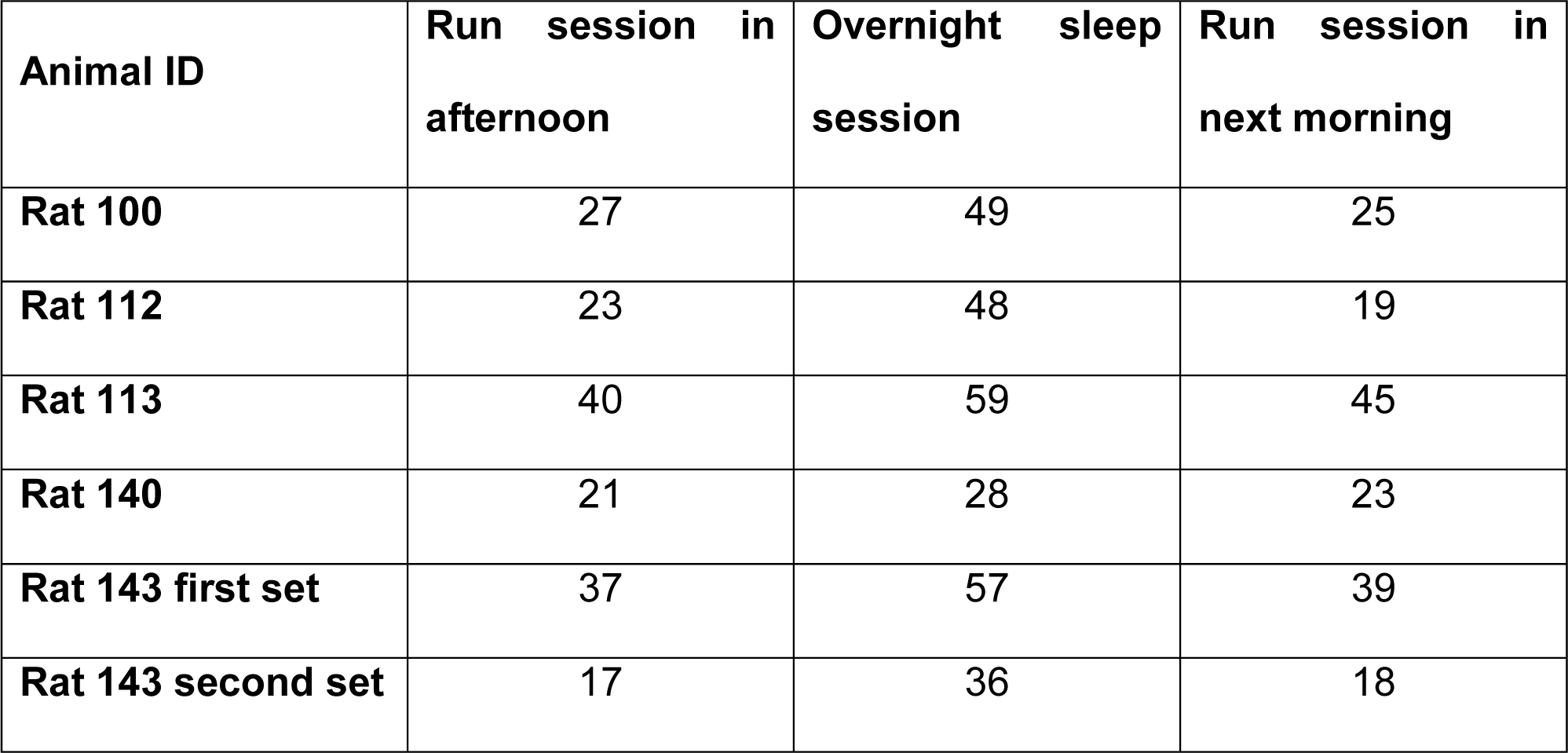
Number of simultaneously recorded CA3 cells in each recording session

**Table 2.** Number of CA3 cells in each cell group from each overnight sleep session

| **Cell group** | **Rat 100** | **Rat 112** | **Rat 113** | **Rat 140** | **Rat 143 first set** | **Rat 143 second set** |
| --- | --- | --- | --- | --- | --- | --- |
| **Novel** | 9 | 1 | 5 | 2 | 11 | 4 |
| **Familiar** | 3 | 2 | 11 | 2 | 5 | 6 |
| **Neither** | 32 | 35 | 33 | 19 | 37 | 24 |
| **Both** | 5 | 10 | 10 | 5 | 4 | 2 |

**Table 3.** Number of CA1 cells in each cell group from each overnight sleep session

| **Cell group** | **Rat 100** | **Rat 112** | **Rat 113** | **Rat 140** | **Rat 143 first set** |
| --- | --- | --- | --- | --- | --- |
| **Novel** | 5 | 0 | 3 | 0 | 0 |
| **Familiar** | 2 | 1 | 2 | 0 | 0 |
| **Neither** | 2 | 9 | 15 | 1 | 1 |
| **Both** | 8 | 3 | 14 | 1 | 3 |

Pearson’s correlations, two-way ANOVAs, and two-way repeated measures ANOVAs were performed using *corr, fitlm,* and *fitrm* functions in Matlab, respectively. Multiple comparisons were performed only when a significant interaction was found between variables included in ANOVAs. The permutation test used for replay analyses shuffled the types of candidate replay events (i.e. Novel or Familiar) 5000 times to obtain a null distribution for variables of interest (see Figures 5B-F). Monte Carlo p-values were calculated using the formula: (N_subset_+1)/(N_shuffle_+1), where N_subset_ is the number of shuffles with values greater than or less than the observed value (two-tailed test) and N_shuffle_ is the total number of shuffles.

**Figure 5.**
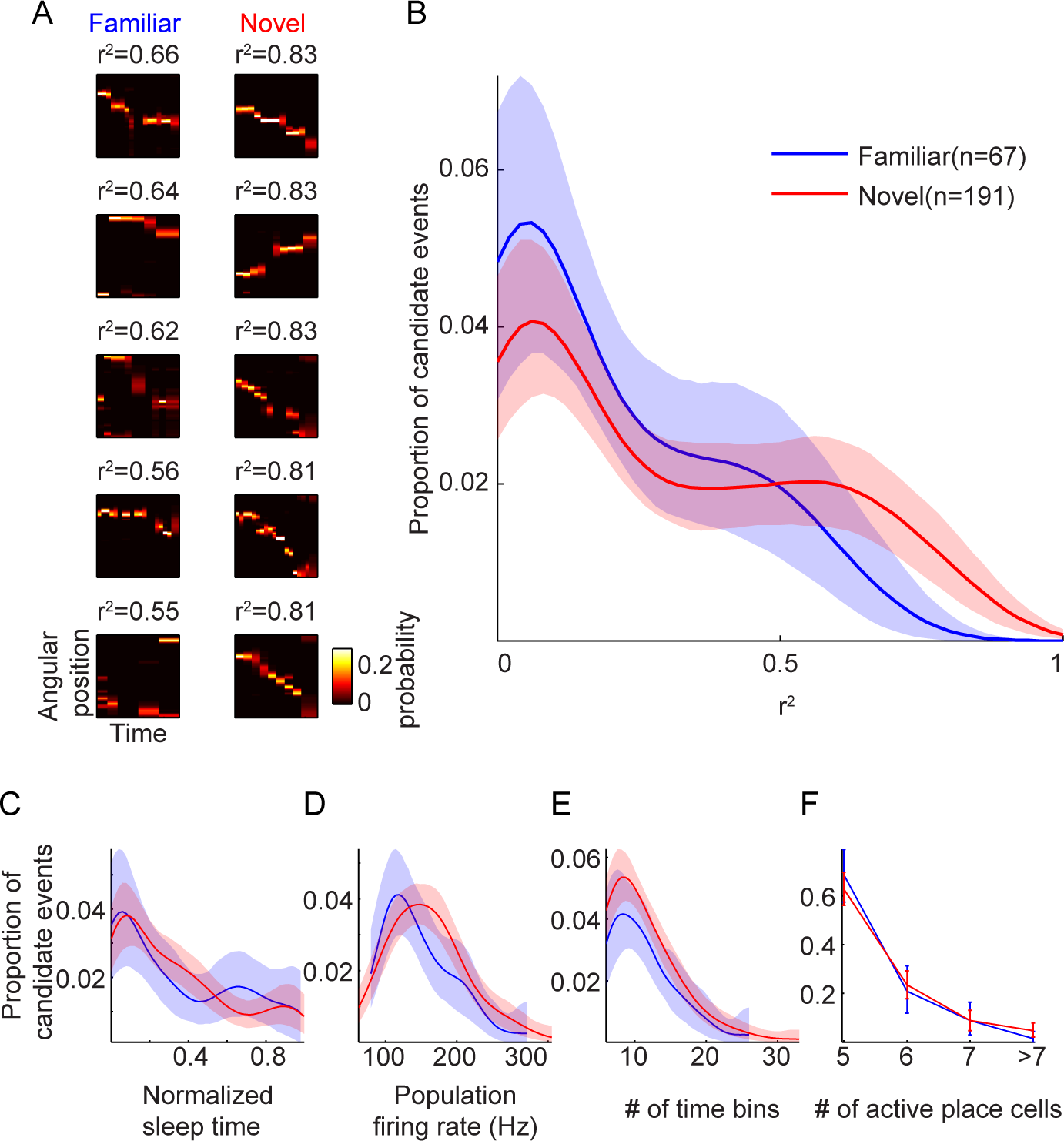
Replay fidelity was higher for CA3 place cell ensembles representing a novel environment than for ensembles representing a familiar environment. ***A***, Candidate replay events with the top five r^2^ values in each group (i.e., Novel and Familiar) are shown. The color scale corresponds to the position probability from Bayesian decoding, with the brightest color representing maximal probability. Circular-linear regressions were computed between angular positions and time to obtain a r^2^ value for each candidate replay event. ***B***, Distributions of r^2^ values for each type of candidate event (i.e., Novel and Familiar) are shown. ***C-F***, Other characteristics of candidate replay events are shown: timing within overnight sleep when a candidate event occurred (C); population firing rate within a candidate event (D); duration, or number of time bins, of a candidate event (E); and number of different CA3 cells that fired at least one action potential within a candidate event (F). Distributions in B-E were obtained using kernel density estimation with a Gaussian kernel (SD = 0.1 x maximum range of values). Shaded regions and error bars indicate 95% confidence intervals.

Variability within our data is shown by 95% confidence intervals. Confidence intervals were computed by bootstrapping using the *bootci* function in Matlab. We used 5000 bootstrapped samples drawn with replacement to estimate each confidence interval.

### Code and data accessibility

Matlab scripts were custom written for the analyses in this paper. Scripts and data are available upon request.

## Results

### CA3 place cell ensemble representations of a novel track emerged across laps

A previous study reported that stable representations of novel spatial environments emerge in CA3 place cell ensembles after approximately 20-30 minutes of experience (Leutgeb et al., 2004). Therefore, we expected that CA3 place cell ensembles would exhibit initially unstable representations of a novel track that would gradually stabilize across multiple exposures. To confirm this, we examined the dynamics of position tuning from CA3 place cell ensembles across successive laps on a novel circular track and compared results to CA3 place cell firing across laps on a familiar track. As is apparent in an example recording from one rat, firing patterns of CA3 place cells remained stable across laps on the familiar circular track but changed across laps on the novel circular track (Figure 2A). We used a population vector analysis (Leutgeb et al., 2005) to quantify the change in place cell ensemble activity between pairs of laps on the familiar or the novel circular track. For each lap on the track, firing rates from all simultaneously recorded CA3 place cells were combined (see Materials and Methods). We then assessed the similarity of CA3 place cell population activity for a pair of laps as a function of lap number difference. The degree of familiarity of the track (i.e., novel or familiar) significantly affected how much ensemble activity correlations decreased as the lap number difference increased. In general, ensemble activity across laps on the familiar track was significantly more similar than was ensemble activity across laps on the novel track (Main effect of track type in multiple regression analysis: β = −0.039, F(1,4372) = 1180, p = 1.93×10^−229^). Moreover, ensemble activity correlations on the novel and familiar tracks were differentially affected by the amount of time between laps (Figure 2B; interaction between track type and lap number difference: β = −0.0041, F(1,4372) = 350, p = 3.71×10^−75^). Correlation values for the familiar track decreased slightly as time between laps increased (Pearson’s r = −0.0502, p = 0.0057), whereas correlation values for novel lap pairs decreased rapidly as time between laps increased (Pearson’s r = −0.448, p = 2.47×10^−67^). These results confirm that CA3 place cell ensemble activity patterns change in a novel environment more than in a familiar one, suggesting that plasticity occurs in the CA3 network during learning of a novel environment.

### CA3 place cells that represented a novel track were selectively reactivated during SWRs in NREM sleep

Presumably, newly formed place cell representations of novel places must be consolidated so that memories of these places can be stably retrieved in the future. As explained above, memory consolidation is thought to occur during NREM sleep. Thus, we examined whether novelty-dependent changes in CA3 place cell ensemble activity during active awake behaviors persisted into subsequent NREM sleep. To assess novelty-dependent effects, we separated CA3 place cells into different categories according to whether they were active on a novel track, a familiar track, both, or neither (i.e., Novel, Familiar, Both, and Neither groups; see Materials and Methods). In previous studies, CA1 place cells that represented novel locations during awake behaviors were more likely to be reactivated during high frequency ripples (Cheng and Frank, 2008), and CA3 and CA1 place cells that represented the same novel locations were shown to reactivate simultaneously during subsequent sleep (O’Neill et al., 2008). Thus, we hypothesized that CA3 place cells representing novel environments would be preferentially reactivated during SWRs in subsequent NREM sleep compared to CA3 place cells representing familiar environments. Indeed, we found that CA3 place cells that were active on a novel track (n = 32) showed firing rate increases near the onset of ripples in subsequent NREM sleep that were significantly greater than firing rate increases for place cells that were active on a familiar track (n = 29) (Figure 3A, C; interaction between group and time since ripple onset: two-way repeated measures ANOVA, F(3,273) = 13.9, p = 1.75×10^−8^; Tukey post-hoc test, Novel vs. Familiar: p = 1.69×10^−4^). Firing rates of place cells increased to a similar extent in novel and familiar groups during ripples recorded on the night before exposure to the novel environment (Figure 3B, D; interaction between group and time since ripple onset: two-way repeated measures ANOVA, F(1,55) = 3.62, p = 0.0624).

However, the analyses above did not consider place cells that were active in both novel and familiar environments but showed preferential activation for one environment or the other (i.e., CA3 place cells in the Both category, which showed a range of preferential firing selectivity for the novel and familiar tracks). Thus, we implemented a selectivity index to quantify on a numerical scale how selective each CA3 place cell’s firing was for the novel or familiar track (see Materials and Methods). We found that peak normalized firing rates within SWRs were positively correlated with selectivity for the novel environment only for SWRs recorded the night following the novel experience but not for SWRs recorded the night before the novel experience (Figure 3E; significant interaction between selectivity index and night: F(1,87) = 11.5, p = 0.00104; significant correlation between selectivity index and peak normalized firing rate for night after novel experience: r = 0.419, p = 4.32×10^−5^; no significant correlation between selectivity index and peak normalized firing rate for night before novel experience: r = 0.132, p = 0.217). Consistent correlation results were observed when firing rates were normalized using a different baseline (Supplementary Figure 2). These results suggest that CA3 place cells that preferentially coded a novel environment were more strongly reactivated during SWRs in subsequent NREM sleep than were CA3 place cells that preferentially represented a familiar environment.

### CA3 place cells preferentially replayed representations of recently learned trajectories during SWRs in NREM sleep

The enhanced firing of CA3 place cells representing the novel track during SWRs raises the possibility that CA3 place cell ensembles preferentially replay novel trajectories during SWRs in NREM sleep. To test this hypothesis, we first set out to establish the viability of assessing replay in our overnight recordings by determining whether CA3 place cell representations remained stable across overnight sleep. To assess the extent to which CA3 place cell representations of familiar and novel trajectories remained stable across overnight sleep recordings, we used the position firing rate maps before sleep to reconstruct rats’ positions from ensemble spiking activity on the circular track after sleep using a probabilistic approach based on Bayes’ rule (Zhang et al., 1998). An example Bayesian decoded spatial probability distribution that corresponded to a lap on the circular track in the novel room is shown in Figure 4A. The peak of the decoded probability distribution closely followed the rat’s actual position, especially when the rat was moving. Decoding accuracy using this approach is shown for each rat in the novel (Figure 4B) and familiar (Figure 4C) environments. Most rats’ recordings (Rats 100, 112, 113, and 143 in the novel room and Rats 112, 113, and 143 in the familiar room) surpassed our criterion for sufficient decoding accuracy (see Materials and Methods) and were thus included in subsequent replay analysis during NREM sleep.

To evaluate the extent to which familiar or novel experiences were replayed during NREM sleep, we estimated spatial representations coded by CA3 place cell ensembles during candidate replay events using Bayesian decoding (see Materials and Methods). Candidate events containing ensembles active on the novel or the familiar track were categorized as Novel and Familiar groups, respectively. Following standards set by previous studies, we assessed the fidelity of replay by fitting a line to the decoded distributions across time and then using corresponding r^2^ values to represent the fidelity of replay (Davidson et al., 2009; Karlsson and Frank, 2009). r^2^ values closer to 1 correspond to higher fidelity replay. The candidate events with the top 5 r^2^ values for Novel and Familiar groups are shown in Figure 5A. As can be seen in these selected events, reconstructed positions for the Novel group resembled novel trajectories better than reconstructed positions for the Familiar group resembled familiar trajectories. The distributions of r^2^ values across all candidate replay events for novel and familiar environments are shown in Figure 5B. Overall, the r^2^ values for novel replay events (n=191) were significantly higher than r^2^ values for familiar replay events (n=67) (Permutation test, 5000 shuffles, p = 0.0018). However, the distributions of r^2^ values may have been affected by several variables associated with candidate events that are unrelated to novelty per se. We thus assessed whether differences in other characteristics of the candidate events (i.e., timing within overnight sleep, population firing rate, duration of candidate events, and number of active place cells) were observed between novel and familiar replay events. No significant differences between novel and familiar groups were observed for any of these measures (Figure 5C-F; Permutation test, 5000 shuffles, Normalized sleep time: p = 0.565, Population firing rate: p = 0.0832, Duration: p = 0.773, Number of active place cells: p = 0.210). These results suggest that ensembles of CA3 place cells preferentially replay novel experiences during NREM sleep.

### CA3 place cells reduced their firing rates during REM sleep

A previous study reported that putative pyramidal cells in CA1 increase their firing rates during NREM and decrease their firing rates during REM sleep, suggesting that REM may serve to prevent accumulating increases in firing rates across successive NREM episodes (Grosmark et al., 2012). Another related study reported that firing rates of CA3 putative pyramidal cells reduced their firing rates more in REM sleep relative to NREM sleep compared to CA1 cells (Mizuseki et al., 2012). These findings, taken together with the increased firing rates during SWRs that we observed for CA3 place cells preferentially representing the novel track (Figure 3), suggest that particularly strong firing rate decreases may occur during REM for CA3 place cells that preferentially code novel environments. To investigate whether novelty-dependent reductions in CA3 firing rates occurred during REM, we compared normalized firing rates of CA3 place cells with fields on only the novel track or only the familiar track across NREM and REM episodes. Both groups of CA3 place cells (i.e., Novel and Familiar) decreased their firing rates from NREM to REM periods of sleep on the night preceding the novel experience and the night following the novel experience (Figure 6A-D; Main effect of normalized time: two-way repeated measures ANOVA, F(1,273) = 2.44×10^3^, p = 3.99×10^−139^; no significant interaction between neuronal group and normalized time, F(3,273) = 0.612, p = 0.608). This pattern of results suggests that CA3 place cell firing rates decrease during REM regardless of novel experience. However, a small effect of novel experience on firing rate decreases during REM was observed when all CA3 place cells were included and categorized according to their selectivity for the novel or the familiar environment. CA3 place cells showed slightly greater firing rate reductions during REM as their preference for coding the novel track increased, and this correlation was only observed during REM sleep recordings after the novel experience (Figure 6E; significant interaction between selectivity index and night: F(1,87) = 4.68, p = 0.0333; significant correlation between selectivity index and minimum normalized firing rate for night after novel experience: r = −0.246, p = 0.0202; no significant correlation between selectivity index and minimum normalized firing rate for night before novel experience: r = −0.0093, p = 0.931). A previous study showed that firing rate decreases over sleep correlate with the rate of SWRs in NREM sleep (Miyawaki and Diba, 2016). However, the weak relationship between preferential coding of the novel environment and CA3 place cell firing rate decreases during sleep after the novel experience is unlikely to be explained by this effect considering that the rates of SWRs did not differ during sleep preceding and following the novel experience (Supplementary Figure 3). Moreover, firing rates rebounded to pre-REM levels during the next NREM period (Figure 6A-D). Whether such weak and transient effects would significantly impact processing of novel memories during sleep is unclear.

**Figure 6.**
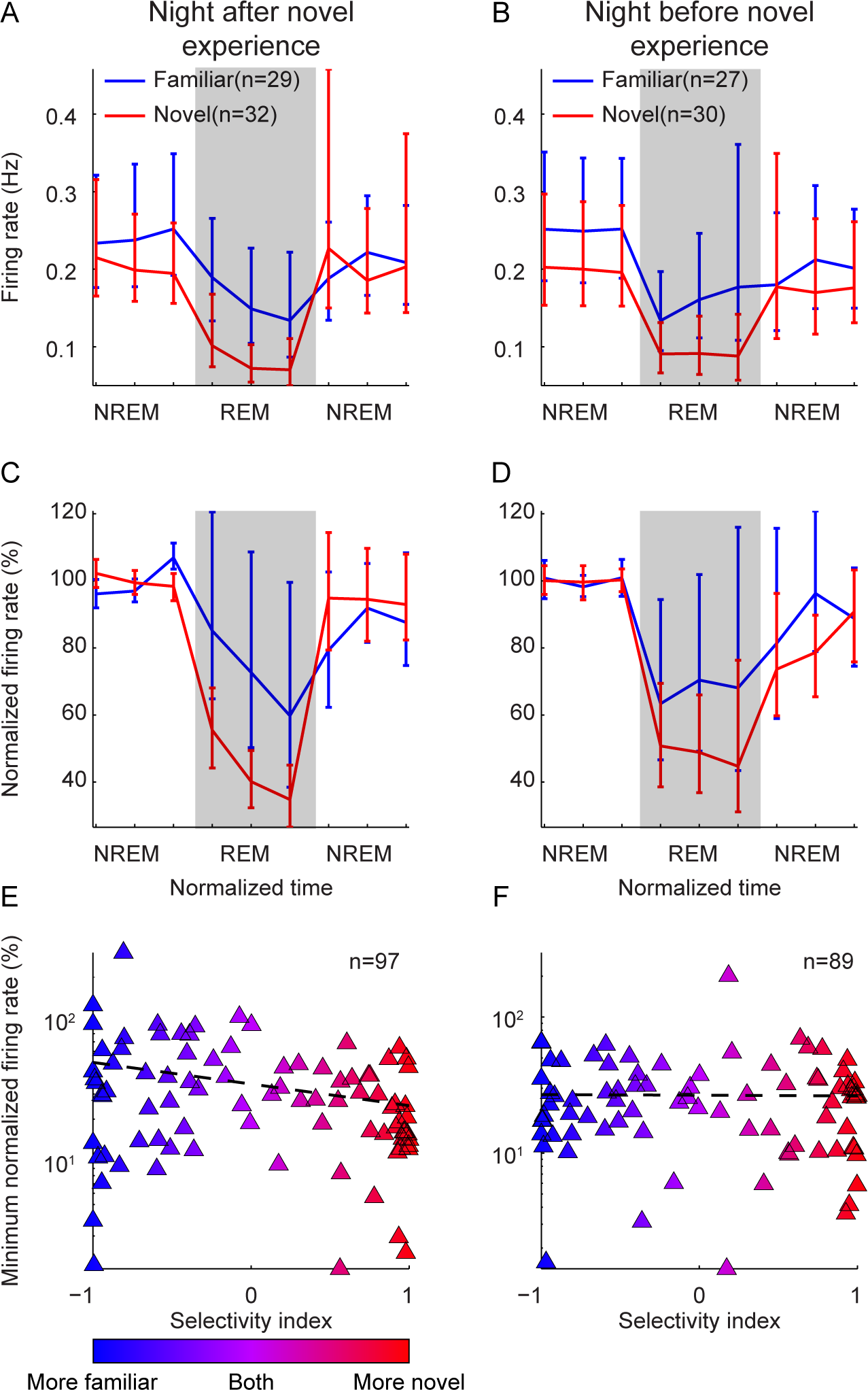
CA3 place cells decreased their firing rates during REM sleep. ***A-B***, Firing rates of CA3 cells decreased from NREM to REM during sleep following (A) and preceding (B) a novel experience. The firing rate for each CA3 cell was averaged across the sleep episodes that contained REM periods. Error bars indicate 95% confidence intervals. ***C-D***, Normalized firing rates of CA3 place cells in Novel and Familiar groups decreased similarly from NREM to REM during sleep following (C) and preceding (D) a novel experience. The firing rate for each CA3 cell was normalized by the mean firing rate in the NREM period that immediately preceded the REM period. Durations of NREM and REM periods in sleep episodes were normalized (see Materials and Methods). ***E-F***, Individual place cells’ minimum normalized firing rates during REM were negatively correlated with the selectivity index described in Figure 3 during sleep following (E) but not preceding (F) a novel experience. The black dashed line shows the linear fit to the data.

## Discussion

A major question in sleep research is how new memories are consolidated during overnight sleep. Memories involving the hippocampus are thought to be initially stored in the CA3 recurrent network (Marr, 1971; Treves and Rolls, 1992; Steffenach et al., 2002), and CA3 is required for consolidation of hippocampal-dependent memory (Nakashiba et al., 2009). Yet, most in vivo studies of hippocampal neuronal activity during active exploratory behaviors (i.e., when memories are formed) and subsequent sleep (i.e., when recently formed memories are thought to be consolidated) have focused on CA1 (Pavlides and Winson, 1989; Wilson and McNaughton, 1994; Kudrimoti et al., 1999; Nádasdy et al., 1999; Lee and Wilson, 2002). Although previous studies have shown that CA1 place cell reactivation during SWRs is stronger for cells representing novel places compared to familiar places (Cheng and Frank, 2008; O’Neill et al., 2008), supporting a role for SWRs in the consolidation of new memories, analogous studies for CA3 place cells were lacking. To address this disparity, we investigated changes in CA3 place cell firing patterns during overnight sleep after experiences in both novel and familiar environments. We found that CA3 place cells that represented novel environments were preferentially recruited during SWRs (Figure 3) and that CA3 place cell ensembles replayed representations of novel environments during NREM sleep with higher fidelity than representations of familiar environments (Figure 5).

The present results imply that memory consolidation during sleep may be an active process that selects newly encoded information rather than a passive process that reflects the duration of recent experiences. If the latter were the case, one would not expect preferential reactivation of CA3 representations of novel experiences during sleep since rats spent equal amounts of time in familiar and novel environments before sleep. Although rats did not perform a memory task to directly demonstrate how well they remembered the novel environment after sleep, CA3 place cell ensemble representations of novel environments remained stable after sleep (Figure 4), suggesting successful retrieval of spatial representations that were formed before sleep. Also, previous studies have shown that disrupting SWR-related neuronal activity during NREM sleep impairs memory performance (Girardeau et al., 2009; Ego-Stengel and Wilson, 2010) and corrupts spatial representations of a novel, but not a familiar, environment (van de Ven et al., 2016). Therefore, a plausible hypothesis is that novel information is preferentially consolidated during SWRs in part due to enhanced reactivation of CA3 cells that encode novel experiences (Figure 3). Together, these results suggest a central role for SWRs in consolidation of new memories.

However, the synaptic mechanisms that allow SWRs during NREM sleep to preferentially reactivate cells that coded novel experiences remain unknown. There are at least two possible ways to selectively increase the relative strength of synaptic connections underlying novel memory storage. A direct way is to strengthen synaptic connections between neurons that encode novel information. In support of this idea, an *in vivo* study showed that stimulation of CA1 putative pyramidal cells during a transient period consisting of 250 detected SWRs increased neuronal activity during subsequent SWRs (King et al., 1999). Furthermore, stimulating CA1 pyramidal cells and Schaffer collaterals *in vitro* using stimulation protocols patterned after CA3 and CA1 place cell firing during SWRs in vivo has been shown to potentiate CA1 cells’ responses to Schaffer collateral stimulation (Sadowski et al., 2016). Thus, preferential recruitment of CA3 place cells during SWRs that coded earlier novel experiences may selectively strengthen their synaptic connections. On the other hand, a second possible way to selectively increase the relative strength of synaptic connections underlying newly encoded memories is to weaken synaptic connections between neurons that do not store novel information. Several studies have suggested that SWRs can weaken synaptic strength (Colgin et al., 2004; Bukalo et al., 2013; Norimoto et al., 2018). In particular, a recent study reported that silencing SWR-related neuronal activity during NREM sleep prevents spontaneous weakening of synapses and impairs subsequent learning of new memories (Norimoto et al., 2018). This study further showed that place cells active in a novel environment maintained their firing rates during SWRs across NREM sleep while other hippocampal cells’ firing rates gradually declined during SWRs across the course of NREM sleep. Firing rates of cells representing novel environments may be affected by both of these mechanisms. The two potential roles of SWRs in synaptic plasticity are not mutually exclusive as long as strengthening and weakening of synaptic connections occur in different subsets of synapses.

However, although the above hypotheses assume that firing rate changes are related to changes in synaptic strength, it is important to note that firing rate changes do not directly reflect changes in synaptic strength (Cirelli, 2017). The reported effects could also reflect changes in intrinsic excitability, particularly considering that intrinsic excitability in CA1 pyramidal neurons has been shown to be increased by hippocampal-dependent learning (Mckay et al., 2009). Thus, more studies are needed to directly investigate potential changes in intrinsic excitability and synaptic plasticity in CA3 pyramidal neurons following experience in a novel environment.

More studies are also needed to determine the extent to which the preferential firing rate increases, and high quality replay, observed during SWRs for CA3 place cells that preferentially represented novel environments relate to behavioral performance in newly learned memory tasks. The environment used in the present study was a simple circular track with no measurable memory component. A previous study showed that contextual fear conditioning increased firing rates of CA1 neurons during subsequent sleep in mice (Ognjanovski et al., 2014), consistent with the findings reported here. However, no place cells were reported in this previous study. Instead, the cell ensembles in the study by Ognjanovski and colleagues appeared to primarily contain fast-spiking interneurons, based on the reported firing rates (i.e., ∼5-10 Hz mean firing rates in their Figure 3), which were considerably higher than the sparse mean firing rates that are characteristic of place cells (e.g., ∼0.1-1 Hz mean firing rates in Figures 3A and 6A of the present study; see also Jung et al., 1994). Consistent with this assumption, the same group later reported increases in firing rates of fast-spiking interneurons during SWRs following contextual fear conditioning (Ognjanovski et al., 2017) and furthermore linked changes in interneuron activity during NREM to consolidation of contextual fear memory (Ognjanovski et al., 2018). However, the recordings in the present study were targeted toward place cells and did not contain fast-spiking interneurons. Thus, how the present place cell results relate to previous results involving firing rate changes in fast-spiking interneurons during sleep remains a question for future study.

When comparing the present results in CA3 to previous results in CA1, discussed above, it is important to keep in mind that many differences exist between subregions. Place cell ensemble activity develops more slowly in CA3 than in CA1 after exposure to a novel environment (Leutgeb et al., 2004). Also, in a study that repeatedly exposed rats to an initially novel room, the population firing rate of CA3 cells remained consistent while the CA1 population firing rate gradually declined across multiple days of exposure (Karlsson and Frank, 2008). Moreover, during learning of new goal locations, many CA1, but not CA3, place cells shifted their firing fields to new goal locations (Dupret et al., 2010). These differences between CA3 and CA1 raise the question of how exactly novel memories would be stored in synapses in the two subregions and the extent to which changes in CA3 drive changes in CA1. These remain questions for future study, as the CA1 place cell yield in the present study was too low to permit comparisons across subregions.

Another open question relates to how memories represented in CA3 and CA1 are consolidated. The reactivation of memory representations during sleep is traditionally thought to promote consolidation of memories in neocortex (Squire and Alvarez, 1995). However, while CA1 projects directly to extrinsic cortical areas, including the retrosplenial cortex and the medial prefrontal cortex, CA3 does not (Cappaert et al., 2015). Thus, such a consolidation process would be expected to rely on CA1, with consolidation of memory representations stored in CA3 requiring transmission through CA1. Yet, the extent to which CA3 and CA1 place cell ensembles replay representations of the same sequences of spatial locations at the same time remains largely unknown. In summary, much work remains to be done to understand how replay of representations of novel experiences by place cells in CA3 and CA1 relates to consolidation of spatial memories.

## Supplementary Figure Legends

**Supplementary Figure 1.**
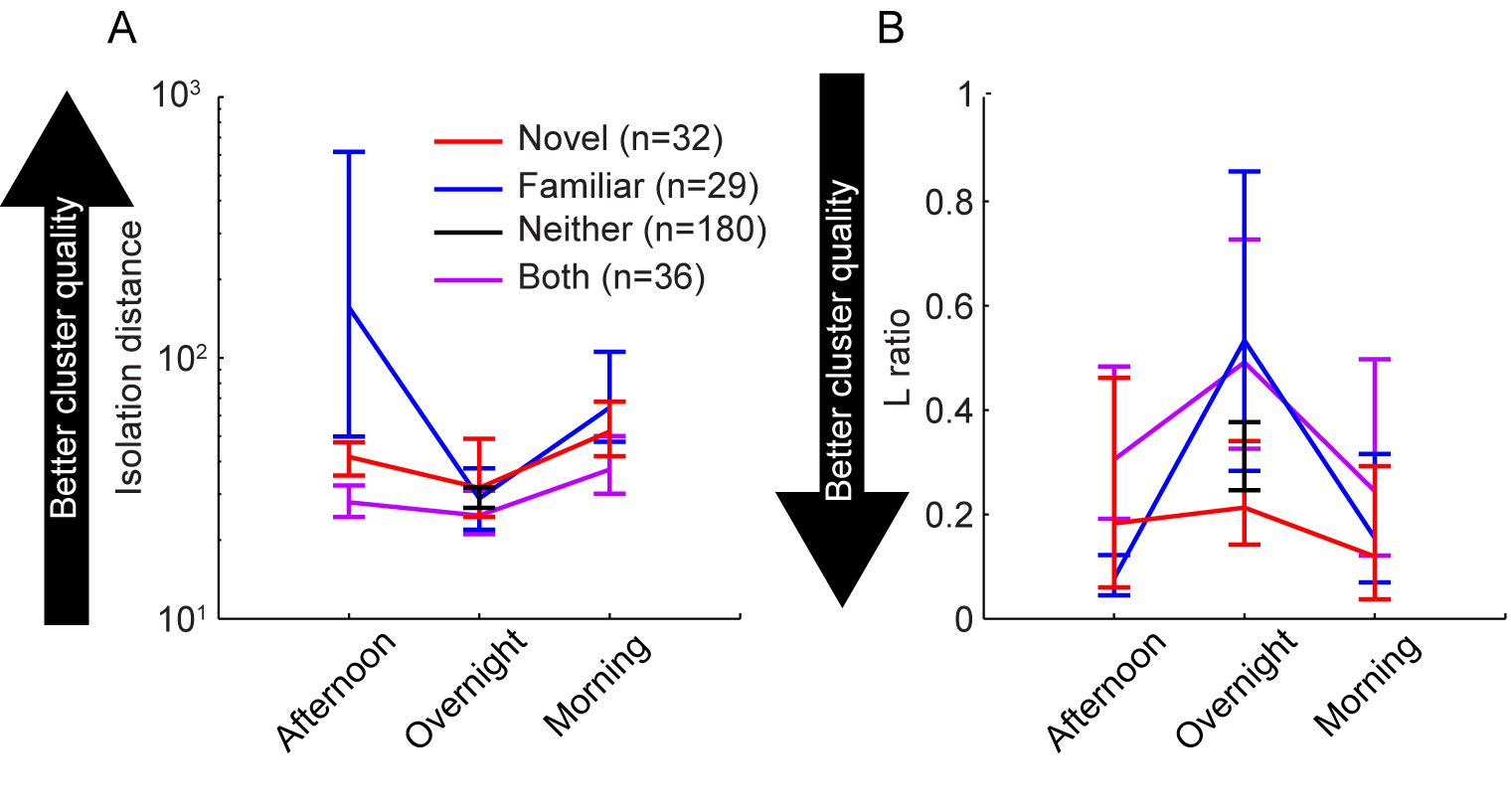
Cluster quality of CA3 units across recording sessions. ***A,*** Isolation distance (higher values indicate better cluster quality) did not differ across different groups of CA3 units (see Materials and Methods for group definitions). ***B,*** L ratios (lower values correspond to better cluster quality) did not differ across different groups of CA3 units.

**Supplementary Figure 2.**
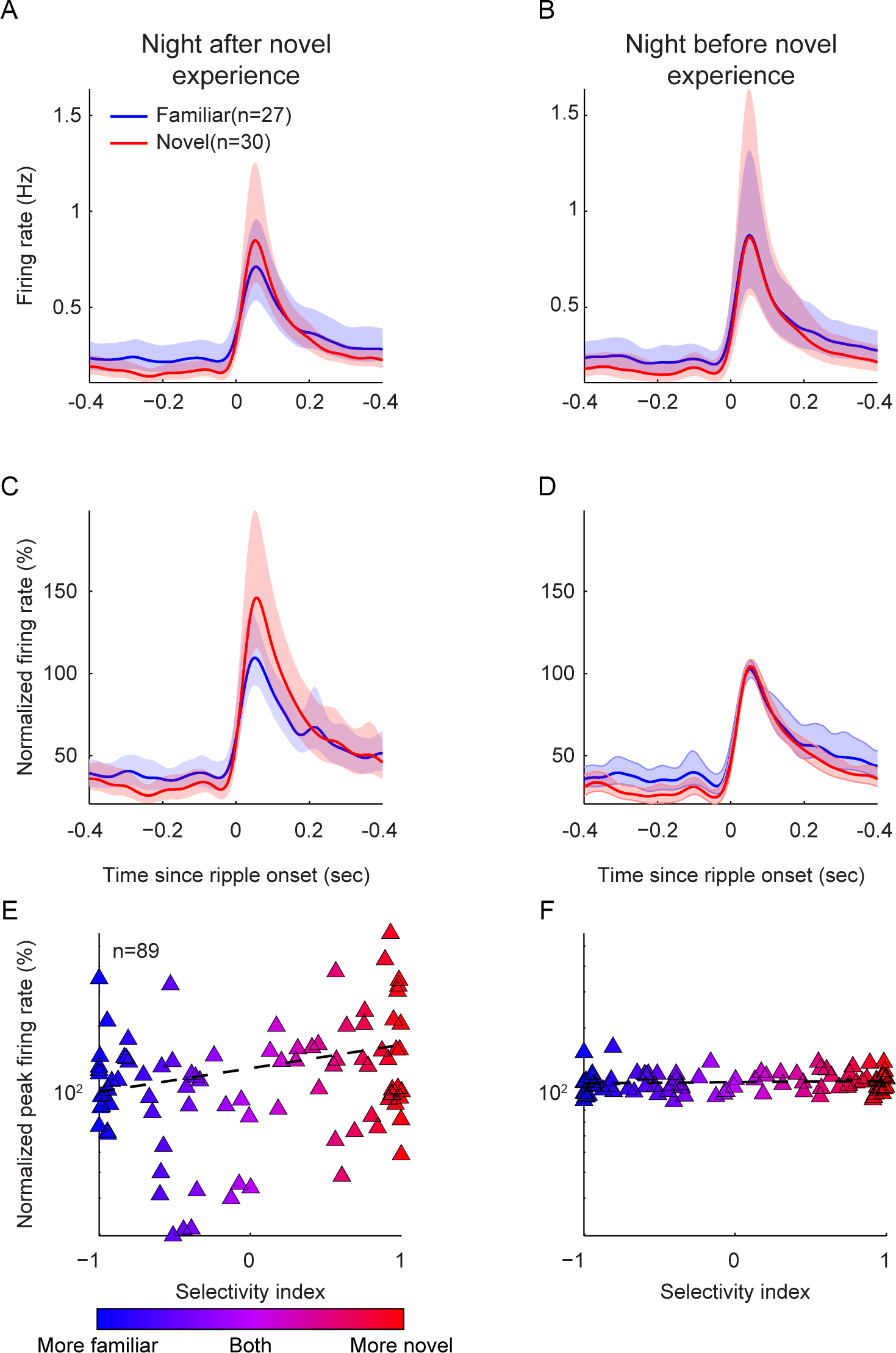
Comparison of firing rates during SWRs for CA3 place cells that preferentially represented novel or familiar environments. In this figure, firing rates were normalized according to firing rates during ripples recorded during sleep prior to the novel experience. Only CA3 place cells that remained stable and thus could be tracked across two consecutive overnight recordings were included in this figure, which is why cell counts are slightly lower than those in Figure 3. ***A-B***, CA3 place cell mean firing rates increased during SWRs recorded during sleep following (A) and preceding (B) a novel experience. Firing rates for each CA3 place cell were averaged across detected SWRs. Shaded areas represent 95% confidence intervals. ***C-D***, Normalized in-ripple firing rates of CA3 place cells in the Novel group increased to a similar extent as CA3 place cells in the Familiar group during sleep following (C) and preceding (D) a novel experience. ***E-F***, Peak normalized firing rates during SWRs were positively correlated with a selectivity index during sleep following a novel experience (E) but not during sleep preceding a novel experience (F) (significant interaction between selectivity index and night: F(1,87) = 6.72, p = 0.0112; significant correlation between selectivity index and normalized peak firing rate for night after novel experience: r = 0.269, p =0.0109 in E; no significant correlation between selectivity index and normalized peak firing rate for night before novel experience: r = 0.0793, p = 0.460 in F). The selectivity index quantified how selective activity of a CA3 place cell was for the familiar room (selectivity index = −1 for maximal selectivity for the familiar room) or the novel room (selectivity index = 1 for maximal selectivity for the novel room). The black dashed line shows the linear fit to the data.

**Supplementary Figure 3.**
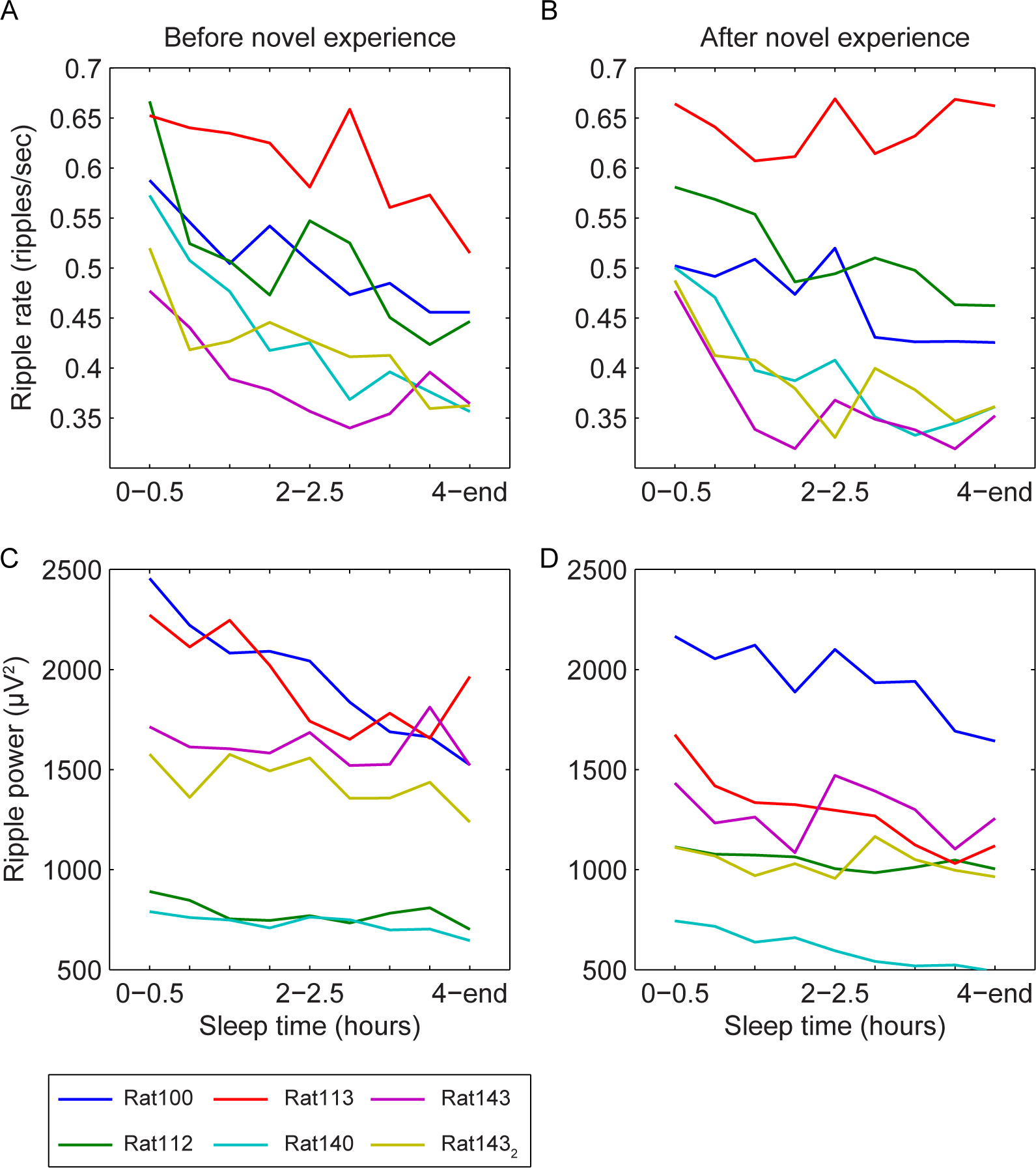
Changes in ripple rate and power across overnight sleep. ***A-B***, Rates of ripple events declined across overnight sleep before (A) and after (B) rats experienced a novel track in a novel room (Main effect of time: F(8,72) = 4.08, p = 4.79×10^−4^). Methods used to detect ripples are described in Materials and Methods (i.e., in the “Identifying SWRs in NREM sleep” section). ***C-D***, Ripple power also declined across overnight sleep before (C) and after (D) novel experience (Main effect of time: F(8,72) = 7.08, p = 7.28×10^−7^). Ripple power was estimated by squaring the amplitude of the ripple-filtered signal (150-250 Hz). Differently colors of lines correspond to measures obtained from overnight sleep recordings from individual rats.

## Acknowledgements

This research was supported by UT Center for Learning and Memory Training Grant #5T32MH106454-03 (E.H.) and National Science Foundation CAREER Award #1453756 (L.L.C.). We thank Devin Wehle, Carlos G. Orozco, and Shelby Brizzolara-Dove for scoring sleep videos; Ayomide Akinsooto, Andrew Wright, and Kayli Kallina for building recording drives and performing histology; Drs. Alexandra Mably, Angel Lopez, John Trimper, Chenguang Zheng, Brian Gereke, and Sean Trettel for insightful discussions and comments. The authors acknowledge the Texas Advanced Computing Center (TACC) at The University of Texas at Austin for providing data storage resources that have contributed to the research described within this paper. URL: http://www.tacc.utexas.edu. The authors declare no competing financial interests.

